# Meta-omic profiling reveals ubiquity of genes encoding for the high value nitrogen rich biopolymer cyanophycin in activated sludge microbiomes

**DOI:** 10.1101/2023.08.22.553871

**Authors:** McKenna Farmer, Rashmi Raj, Will Tarpeh, Keith Tyo, George Wells

## Abstract

Recovering nitrogen (N) from municipal wastewater is a promising approach to prevent nutrient pollution, reduce energy use, and transition towards a circular N bioeconomy, but remains a technologically challenging endeavor. Existing N recovery techniques are optimized for high-strength, low-volume wastewater. Therefore, developing methods to concentrate dilute N from mainstream wastewater will bridge the gap between existing technologies and practical implementation. The N-rich biopolymer cyanophycin is a promising candidate for N bioconcentration and recovery due to its solubility characteristics and potential for high levels of accumulation in a limited number of bacterial isolates. However, the cyanophycin synthesis pathway is poorly explored in natural and engineered microbiomes. In this study, we analyzed over 3700 publicly available metagenome assembled genomes (MAGs) and found that the cyanophycin synthesis gene *cphA* was ubiquitous across common activated sludge bacteria. Surprisingly, we found that *cphA* was present in all analyzed genomes of the common phosphorus accumulating organisms (PAO) *Ca*. ‘Accumulibacter’ and *Tetrasphaera*, suggesting potential for simultaneous N and P bioconcentration in the same organisms. Using metatranscriptomic data, we also confirmed the expression of *cphA* in lab-scale bioreactors enriched with PAO. Our findings suggest that cyanophycin synthesis is a ubiquitous metabolic pathway in activated sludge microbiomes and therefore may have potential for integration in existing biological nutrient removal and recovery processes. We anticipate this work to be a starting point for future evaluations of combined N and P bioaccumulation, with the ultimate goal of advancing widespread adoption of nutrient recovery from municipal wastewater.

## 1. Introduction

Recovering nitrogen (N) from municipal wastewater is a promising method to circularize anthropogenic N use. Traditionally, fertilizer manufacturing and other industries synthetically fix N from the atmosphere through the energy intensive Haber Bosch process, which accounts for 1-2% of global energy use (Batstone et al., 2015). A significant portion of this reactive N is ultimately lost to municipal and industrial wastewater or agricultural drainage water. This reactive N causes harm to aquatic environments and impacts public health (Galloway et al., 2008), therefore reducing N emissions to the environment is therefore a key goal of wastewater treatment plants. Reactive N is typically removed from municipal wastewater through microbially driven redox reactions back to inert N_2_, which is then dissipated back to the atmosphere, rather than recovered. N recovery from wastewater is appealing because it can reduce N emissions to the environment while rerouting reactive N back to food or chemical production, promoting a transition from a linear to a circular anthropogenic N cycle and reducing reliance on Haber Bosch. However, existing N recovery techniques are optimized for high-strength, low-volume wastewater that differ from low-strength, high-volume mainstream municipal wastewater (Beckinghausen et al., 2020). Therefore, a partition-release-recovery (PRR) approach has been proposed to sequester nutrients from mainstream wastewater to a highly concentrated sidestream (Batstone et al., 2015). An efficient partition step to concentrate dilute N is an essential yet poorly explored element of this approach for N recovery.

A promising lead for N bioconcentration is cyanophycin, a bacterial intracellular biopolymer composed of amino acids aspartate and arginine. Cyanophycin is polymerized via cyanophycin synthetase, *cphA* (Ziegler et al., 1998) and broken down into monomers by cyanophycinase, *cphB* (Richter et al., 1999) (Figure S1). Cyanophycin monomers may be hydrolyzed by isoaspartyl peptidase *iaaA*, though other enzymes may also perform this function (Sharon et al., 2023). Cyanophycin was originally characterized in cyanobacteria, and the metabolic pathway has also been identified in a limited number of non-phototrophic bacteria (Füser and Steinbüchel, 2007). In isolate cultures, cyanophycin can reach up to 40% of cell dry weight (Elbahloul et al., 2005), and cyanophycin granules can selectively solubilized by manipulating pH (Füser and Steinbüchel, 2005). Given these attractive properties, industrial biotechnology research has focused on maximizing cyanophycin production in recombinant bacteria, yeasts, and transgenic plants (Du et al., 2019; Nausch et al., 2016).

Cyanophycin production in mixed microbial communities, particularly in wastewater bioprocesses, has not been well-documented to date. Recent work has also shown that cyanophycin can be produced unintentionally in activated sludge (Zou et al., 2022). Other meta-omic studies have incidentally identified cyanophycin synthetase genes in wastewater-associated microbes (Füser and Steinbüchel, 2007; Singleton et al., 2022) but have not performed a specific and systematic search for the cyanophycin pathway in wastewater bioprocesses. Cyanophycin accumulation could add immense value to existing biological nutrient removal practices, namely the enhanced biological phosphorus removal (EBPR) process. EBPR processes enrich phosphorus accumulating organisms (PAO), heterotrophs that release and uptake P under alternating redox conditions and substrate availability. Existing EBPR processes already use the PRR approach for P recovery, where P-rich biomass is bioconcentrated and then physically separated from the dilute liquid stream. Therefore, integrating cyanophycin accumulation with existing P removal practices is attractive, potentially offering a lower barrier to entry for N recovery.

To better understand the potential role of cyanophycin as an N-rich biopolymer for the PRR approach, we assessed over 3700 publicly available metagenome assembled genomes (MAGs) to understand the prevalence and diversity of the cyanophycin biosynthetic pathway in existing wastewater microbiomes. We also curated MAGs and isolate genomes of key functional groups known to contribute to N cycling and P accumulation in wastewater treatment bioreactors to determine whether the associated bacterial taxa (many of which are as-yet-uncultivated) are capable of cyanophycin accumulation. Finally, we analyzed gene expression data of known PAO to understand whether PAO could utilize cyanophycin synthesis genes.

## 2. Methods

### 2.1. Large metagenome assembled genomes datasets

Wastewater MAGs were obtained from two primary sources that used the MIMAG standard to determine MAG quality, where high-quality MAGs met ≥90% completeness and ≤5% contamination and medium-quality MAGs met ≥50% completion and ≤10% contamination (Bowers et al., 2017). The first dataset is Genomes from Earth’s Microbiomes, a dataset assembled by IGM/M Data Consortium from a variety of natural and engineered systems (Nayfach et al., 2021). Out of the entire collection of 52,515 medium and high-quality MAGs, MAGs with the metadata field “ecosystem_category” matching the query “wastewater” were selected for this analysis, resulting in a subset of 2627 MAGs. We annotated the MAGs for coding regions and function using prokka v1.14.6 with default e-value threshold of 1e-06 (Seemann, 2014). The wastewater MAGs from this dataset were primarily represented by anaerobic digester samples, so a second set of MAGs representing activated sludge was also used (NCBI BioProject PRJNA629478). In this study, 1083 high-quality MAGs were recovered through a combination of short-read and long-read sequencing (Singleton et al., 2021).

### 2.2. Wastewater microbe functional groups

Isolate genomes and high-quality MAGs from key wastewater functional groups were downloaded from NCBI. As a point of comparison, cyanobacterial genomes with known cyanophycin metabolism pathways were also included in the analysis. A complete list of genomes used is available in supplemental Table S1. To understand patterns among genes clustered near *cphA*, conserved gene clusters with gene synteny 6000 bp upstream and downstream of *cphA* were identified and grouped using GeneGrouper with default grouping settings (McFarland et al., 2022).

### 2.3. Gene expression

High-quality *Candidatus* ‘Accumulibacter’ (referred to herein as Accumulibacter) MAGs and associated metatranscriptomic data were used to examine *cphA* expression in Accumulibacter PAO. Our group previously operated a lab-scale denitrifying PAO reactor enriched in Accumulibacter (Gao et al., 2017; Wang et al., 2021). Three high quality *Ca*.Accumulibacter MAGs were assembled from this work belonging to clades IA, IC, and IF based on polyphosphate kinase (*ppk1*) gene phylogeny. Metatranscriptomic reads and PAO MAGs metagenomes from this study were accessed through NCBI BioProject PRJNA576469. Raw RNA reads were filtered for quality with fastp (Chen et al., 2018) and reads mapping to rRNA were removed using BBMap (sourceforge.net/projects/bbmap/) against the SILVA database (Quast et al., 2012). Cleaned reads were aligned against Accumulibacter MAGs using kallisto (Bray et al., 2016). Expression levels of each mapped gene were normalized to Transcript per Million Reads (TPM). We also examined MAGs and expression levels from a lab-scale EBPR reactor where *Tetrasphaera* MAGs were recovered (McDaniel et al., 2022). Further operational details of the EBPR reactors can be found in their respective publications.

## 3. Results and Discussion

### 3.1. Cyanophycin metabolism genes are widespread

To understand the relevance of cyanophycin metabolism genes in wastewater microbiomes, we searched for *cphA* in broad collections of wastewater-associated MAGs. The first dataset we examined was primarily represented by MAGs recovered from anaerobic digester sludge (Figure 1). Out of 2627 MAGs, 271 possessed the *cphA* gene, an unexpectedly high prevalence. MAGs from nutrient removal systems and activated sludge were overrepresented in the *cphA* subset compared to their original distribution in the dataset (Figure 1), though the potential overlap between the activated sludge and nutrient removal metadata categories is a limitation of the dataset, since nutrient removal functions are often integrated into activated sludge processes. On the other hand, while nearly three out of four MAGs were derived from anaerobic digesters, these MAGs only represented roughly one out of three MAGs with a *cphA* gene (Figure 1). Examining the MAGs with *cphA* more closely, we found that key nutrient cycling organisms possessed a *cphA* gene, including known PAO Accumulibacter and *Tetrasphaera*, as well as ammonia oxidizing bacteria affiliated with the genus *Nitrosomonas* (Figure S1). This result was surprising, as *cphA* had not been documented in *Ca*. Accumulibacter genomes to our knowledge.

**Figure 1.**
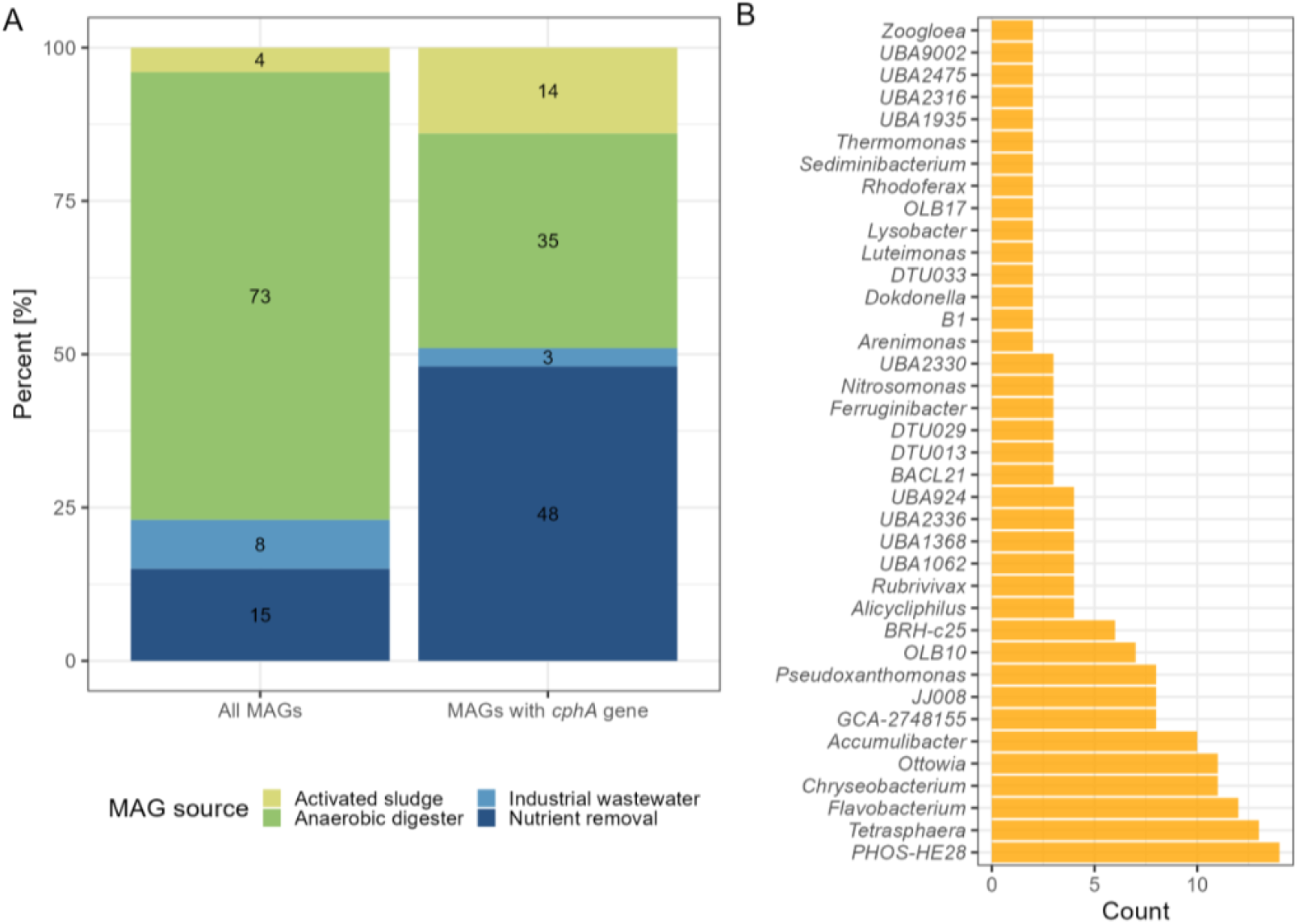
Distribution of MAGs from the Genomes from Earth’s Microbiomes collection (Nayfach et al., 021) based on their identified ecosystem type with percent of total count labelled (A) and genus classifications of MAGs that had a *cphA* gene (B). Data in panel B are shown for genus-level classifications of two or more MAGs based on GTDB taxonomy.

Given the metadata limitations of the GEM dataset, we also looked at the prevalence of *cphA* in a study specifically targeting activated sludge microbiomes performing EBPR and N removal (Singleton et al., 2021). Out of 1083 MAGs from this study, 552 possessed a *cphA* gene copy. Similar to the GEM dataset, we found that N and P cycling microbes harbored the *cphA* genes, including *Ca*. Accumulibacter, *Dechloromonas, Nitrosomonas*, and *Propionivibrio* (Figure S2). We also found that common filamentous bacteria possessed a *cphA* gene copy, including *Zoogloea* and *Kouleothrix*. Although filamentous bacteria are undesirable in large quantities in activated sludge systems due to their contribution to sludge bulking and poor settling, they are ubiquitous throughout activated sludge systems and can improve floc strength in balance with other microbes (Burger et al., 2017). The presence of *cphA* in filamentous bacteria may improve the viability of future cyanophycin applications, as filamentous bacteria can represent over 25% of sludge biomass in well-functioning systems (Araújo Dos Santos et al., 2015; Mielczarek et al., 2012).

A prominent *cphA* harboring genus in both large-scale datasets was PHOS-HE28. These organisms are poorly characterized members of the *Flavobacteriales* order. PHOS-HE28 have been identified in activated sludge and are related to bacteria isolated from saline environments (Bowman, 2020). PHOS-HE28 and other related bacteria possess a *ppk1* gene, so it is possible that these bacteria can store polyphosphate (Lucena et al., 2022), though more examination of *ppk1* phylogeny and other phosphate transport genes is necessary to infer this function. Further investigation of PHOS-HE28 may be a promising avenue for integrating cyanophycin accumulation into existing treatment facilities given its ubiquity in numerous activated sludge microbiomes.

Overall, the unexpectedly high prevalence of MAGs from biological nutrient removal systems with a *cphA* gene is a positive sign that cyanophycin accumulation could integrate with existing nutrient management practices. This finding agrees with recent work that studied cyanophycin gene abundance and production in two full-scale wastewater treatment facilities using biofilm reactors for N and P removal (Zou et al., 2022). They successfully identified *cphA* genes in biomass samples and found an association between *cphA* abundance and Accumulibacter marker gene abundance. However, they used read-based analysis for their metagenomic work rather than assembly-based, so they could not directly associate *cphA* genes with particular taxa. Given the importance of key functional taxa for biological nutrient treatment, we next focused our analysis on specific N and P cycling organisms.

### 3.2. Nitrogen and phosphorus cycling bacteria harbor cyanophycin genes

Since *cphA* genes were widespread among wastewater treatment microbiomes, we next examined a wider suite of complete genomes and near-complete MAGs obtained from NCBI of N and P cycling microbes to determine the potential for cyanophycin accumulation in tandem with existing nutrient removal functions in wastewater treatment bioreactors. We selected genomes of ammonia oxidizing bacteria (AOB), nitrite oxidizing bacteria (NOB), denitrifiers, PAO, and GAO to search for the *cphA* gene. A complete list of genomes examined is available in Table S1.

Out of 68 genomes searched, 34 possessed at least one copy of the *cphA* gene. Notably, nearly all PAO genomes possessed a copy of the *cphA* gene. All Accumulibacter and *Tetrasphaera* genomes had a copy, as well as *Ca*. Dechloromonas phosphoritropha. This finding is consistent with previous research, which identified *cphA* copies in wastewater-associated *Tetrasphaera* (Singleton et al., 2022) and found correlations between Accumulibacter phylogenetic markers and *cphA* gene abundance (Zou et al., 2022). The only analyzed PAO genome without a *cphA* gene copy was a *Ca. ‘*Dechloromonas phosphorivorans’ genome. This result was surprising, particularly since all denitrifying *Dechloromonas* species possessed a *cphA* copy, as did another *Ca. ‘*Dechloromonas phosphorivorans’ genome and the closely related *Ca. ‘*Dechloromonas phosphoritropha’. It is unclear whether this discrepancy is due to a true lack of *cphA* in this particular species or was absent in the MAG due to limitations of sequencing and metagenome assembly. As more high-quality *Ca. ‘*Dechloromonas phosphorivorans’ genomes are assembled and published, the presence of *cphA* in this species will become clearer. Regardless, given that PAO are already harnessed for their affinity for P bioconcentration, the potential for simultaneous N recovery via synthesis of the N-rich biopolymer cyanophycin in the same organism may be promising avenue for combined P and N recovery.

Another notable finding amongst the functional groups was that no NOB genome possessed a *cphA* gene, while multiple AOB genomes possessed a *cphA* gene copy. This finding was surprising since AOB and NOB are similar metabolically as chemolithoautotrophs. Furthermore, some *Nitrospira*-affiliated taxa previously thought to be NOB are capable of complete ammonia oxidation (comammox), resulting in even more metabolic similarities to AOB with the ability to oxidize ammonia (Daims et al., 2016). We analyzed a known comammox genome of *Ca*. Nitrospira nitrosa (van Kessel et al., 2015), and did not find a *cphA* gene. Among the AOB genera, we found *cphA* in all *Nitrosospira* genomes and seven out of 13 *Nitrosomonas* genomes. Of the five *Nitrosomonas* genomes that did not possess *cphA*, four were originally isolated from marine or brackish environments, not activated sludge, and required a salt-enriched medium for growth (Koops et al., 1991). The other *Nitrosomonas* genome that did not possess *cphA* was originally isolated from cattle manure (Nakagawa and Takahashi, 2015). The presence of *cphA* in common wastewater-associated AOB, such as *N. europaea* and *N. nitrosa*, is promising for future study and integration of cyanophycin accumulation with existing nitrification bioprocesses.

We also assessed the presence of *cphB* in all functional group genomes. Interestingly, out of all wastewater-associated genomes, only *Tetrasphaera* species possessed a *cphB* gene, unlike the four cyanobacterial genomes, which all possessed a *cphB* gene copy. The lack of a *cphB* gene does not indicate that a particular organism is incapable of depolymerizing cyanophycin, as other enzymes can depolymerize aspartate-arginine dipeptides such as isoaspartyl peptidase *iaaA*, which can also be employed for other biochemical functions (Sharon et al., 2023). To better understand how cyanophycin could fit into a broader biosynthetic pathway, particularly in genomes lacking a *cphB* gene, we next examined the genes upstream and downstream of *cphA* in each genome.

### 3.3. PAO harbor distinct *cphA* gene clusters

We used GeneGrouper to analyze the genes surrounding *cphA* in each genome and bin the gene clusters into homologous groups. For our gene clustering analysis, we included 4 cyanobacterial genomes and an *Acinetobacter* species with a well-characterized *cphA* gene to understand whether *cphA* gene clusters from as-yet-cultivated taxa from key wastewater treatment functional groups were similar to gene clusters derived from well-studied bacterial isolates.

We found two distinct gene clusters with gene synteny 6000 bp upstream and downstream of *cphA*, shown in Figure 2. Surprisingly, *cphA* gene clusters from *Tetrasphaera* formed their own distinct group (Group 2), while the remainder of the wastewater-associated microbes formed another group (Group 1). None of the cyanobacterial genomes clustered with the wastewater-associated microbes, nor did they form their own homologous cluster.

**Figure 2.**
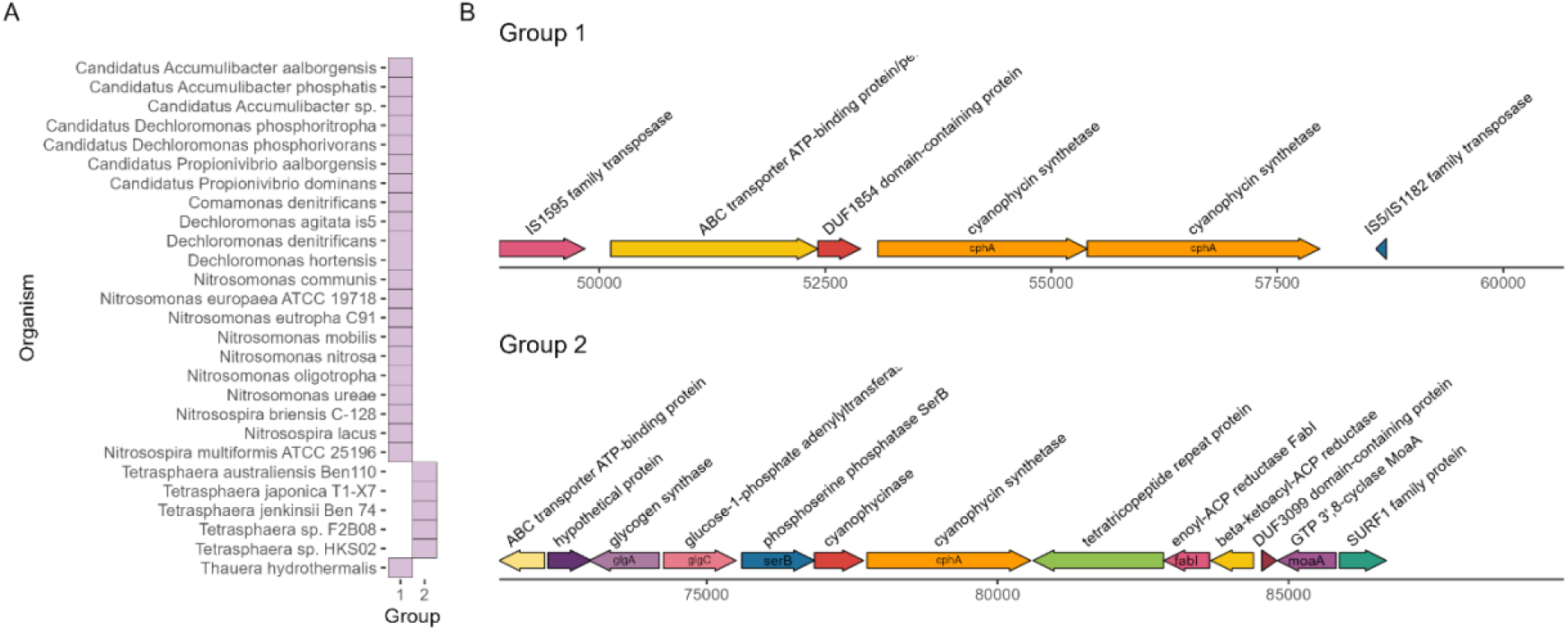
Groups of gene clusters (A) and gene cluster maps (B) around *cphA*.

Another notable finding in the clustering analysis was the complexity of the *Tetrasphaera cphA* gene cluster. The cluster consisted of a copy of *cphB* directly upstream of *cphA*, which is more similar to *cphA* gene clusters of cyanobacteria rather than non-cyanobacterial species (Füser and Steinbüchel, 2007; Krehenbrink et al., 2002). Furthermore, the *Tetrasphaera cphA* gene cluster also consisted of key genes for glycogen synthesis and storage including glycogen synthetase *glgA* and glucose-1-adenylyltransferase *glgC*, as well as the amino acid utilization gene phosphoserine phosphatase *serB*. Glycogen is an important carbon reserve for *Tetrasphaera;* they typically consume glycogen under anaerobic conditions and replenish stores during aerobic conditions (Close et al., 2021; Yu et al., 2023). The clustering of *cphA* and *cphB* with these key glycogen synthesis genes suggests that cyanophycin may be another important carbon and N storage compound for *Tetrasphaera*. Genes that form biosynthetic pathways with increasingly complex metabolites often cluster together (Fischbach et al., 2008; Wolf et al., 2001). More extensive analysis of the *Tetrasphaera* carbon storage metabolism is necessary to understand the potential role of cyanophycin as one member of the pool of carbon storage compounds.

The other *cphA* gene cluster group (Group 1) included a variety of wastewater taxa. This gene cluster consisted of two copies of *cphA*, an unclassified transmembrane transport gene (ATP-binding ABC transporter), and two insertion sequences (IS). The presence of flanking IS indicates that this cluster could be a composite transposon, a type of mobile genetic element that facilitates movement of genetic material within a genome and between bacteria. Flanking IS around functional genes is a hallmark of composite transposons (Siguier et al., 2009). Composite transposons have been studied extensively for facilitating the spread antibiotic resistance and xenobiotic resistance genes via horizontal gene transfer between taxa in diverse microbiomes (Bennett, 2008; Top and Springael, 2003). Further analysis of this *cphA* gene cluster would greatly improve our understanding of whether this is a composite transposon or another type of mobile genetic element, which may have important implications for gene mobilization and transfer in complex, mixed culture microbial communities.

### 3.4. Cyanophycin synthetase is expressed in PAO

Since *Ca*. Accumulibacter would be an excellent candidate for combined N and P bioconcentration, we wanted to determine whether these genes were being expressed *in-situ*. We first examined gene expression by mapping metatranscriptomic reads against three high quality *Ca*. Accumulibacter MAGs assembled from a lab-scale denitrifying P removal bioreactor (Gao et al., 2019; Wang et al., 2021). The samples were obtained over the course of two complete reactor cycles consisting of three redox phases: anaerobic, anoxic (N supplied as nitrite), and aerobic. The reactor was fed with different carbon sources, either acetate or propionate, in equivalent concentrations on a COD basis. The three Accumulibacter MAGs affiliated with different clades — IA, IC, and IF — based on *ppk1* phylogeny and will be referred to hereafter by their clades. All three bins had two neighboring copies of *cphA* present in the genome, which agreed with our previous analysis of Accumulibacter MAGs (section 3.2).

Each of the Accumulibacter MAGs exhibited different expression patterns across redox conditions; bins IA and IF had the greatest *cphA* expression during the aerobic phase, while IC had the greatest *cphA* expression during the anoxic phase (Figure S3). Overall, bin IF had the greatest *cphA* expression, which agrees with previous findings that IF was the most transcriptionally active based on the number of reads mapped to this MAG (Wang et al., 2021). There were no apparent differences in *cphA* expression as a result of different carbon sources, acetate or propionate, in the feed (Figure S3).

In addition to analyzing the *cphA* expression of each MAG, we compared the *cphA* expression to other key functional genes shown in Table 1. We used *glgC, ppk1*, and *phaC* as points of comparison against the expression of *cphA* because these genes are part of important carbon storage and phosphate storage pathways and have been identified in a majority of available *Ca*. Accumulibacter genomes (Petriglieri et al., 2022). The *ppk1* gene in particular is an essential biomarker for PAO activity and is expressed actively in EBPR processes (He et al., 2007; He and McMahon, 2011). As shown in Figure 3, *phaC* had the highest level of expression across all bins. Surprisingly, *cphA* expression was significantly higher than *ppk1* and *glgC* expression in bin IF. While gene expression does not directly indicate microbial activity or kinetics, the level of *cphA* expression relative to *ppk1* and *glgC* points to the possibility of a highly active cyanophycin synthesis pathway.

**Table 1.**
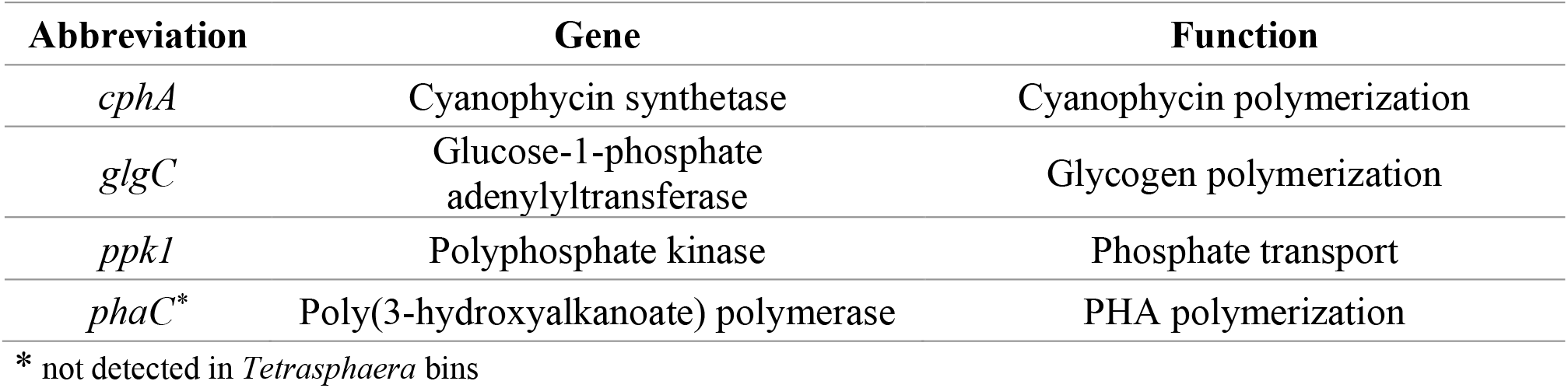
Genes of interest for PAO expression analysis.

| Abbreviation | Gene | Function |
| --- | --- | --- |
| *cphA* | Cyanophycin synthetase | Cyanophycin polymerization |
| *glgC* | Glucose-1-phosphate<br>adenylyltransferase | Glycogen polymerization |
| *ppk1* | Polyphosphate kinase | Phosphate transport |
| *phaC* * | Poly(3-hydroxyalkanoate) polymerase | PHA polymerization |

\* not detected in *Tetrasphaera* bins

**Figure 3.**
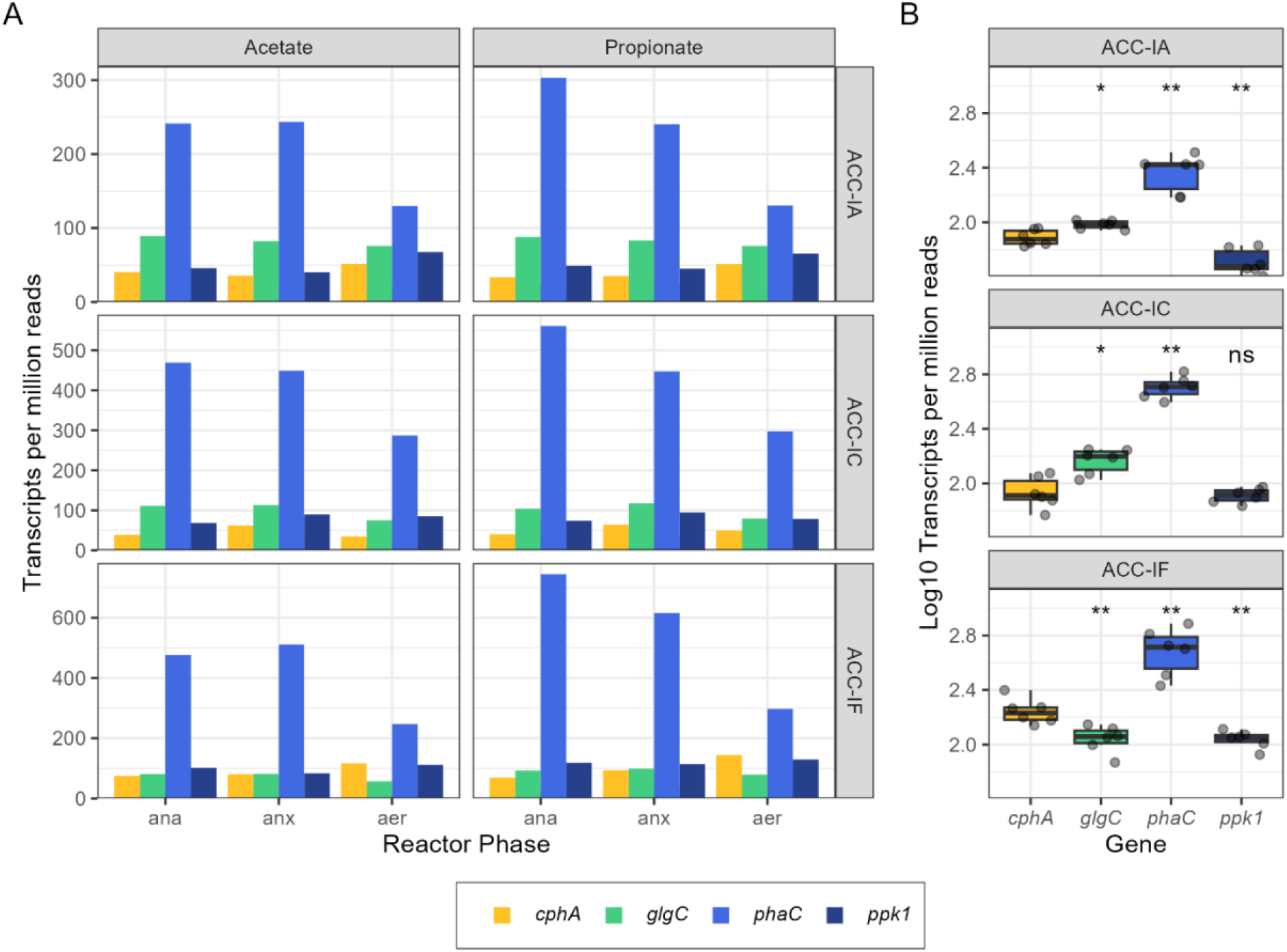
Gene expression profiles from metatranscriptomic reads of *cphA, glgC, phaC*, and *ppk* in *Ca*. Accumulibacter MAGs plotted by the reactor phase (ana=anaerobic, anx=anoxic, aer=aerobic) and carbon source (A) and summarized per MAG (B). Significance levels in panel B are based on the Wilcox rank sum test, where *glgC, phaC*, and *ppk* are compared against *cphA* in each subpanel.

We also examined gene expression data of two *Tetrasphaera* MAGs, TET1 and TET2, recovered from a time-series study of a separate EBPR bioreactor (McDaniel et al., 2022). These MAGs contained two copies of the *cphA* gene in series, similar to *cphA* gene cluster group 1 in Figure 2. On the other hand, TET1 and TET2 had similar genes surrounding *cphA* compared to the other *Tetrasphaera* MAGs in gene cluster group 2 examined in section 3.2, with a neighboring *cphB* as well as other carbon storage and cycling genes (Figure S4). Furthermore, the *cphA* copies in TET1 and TET2 were not surrounded by flanking IS.

Unlike the *cphA* gene copy expression in *Ca*. Accumulibacter, where both copies were expressed evenly, there was a large gap in gene expression between the two *cphA* copies in the *Tetrasphaera* MAGs (Figure S5). Notably, in both MAGs, the longer copy of *cphA* exhibited higher expression levels than the shorter copy; since the gene length is included in the normalization technique of calculating TPM, the gene length should not impact the reported expression levels. The combined gene expression of both *cphA* copies was relatively close to that of *glgC* and *ppk1* in the *Tetrasphaera* bins across the sampling period (Figure 4). Again, gene expression does not directly indicate degree of function or activity, but the comparable level of expression of *cphA* compared to well-understood essential genes of the PAO phenotype is a promising indication that cyanophycin is an active storage polymer for *Tetrasphaera*.

**Figure 4.**
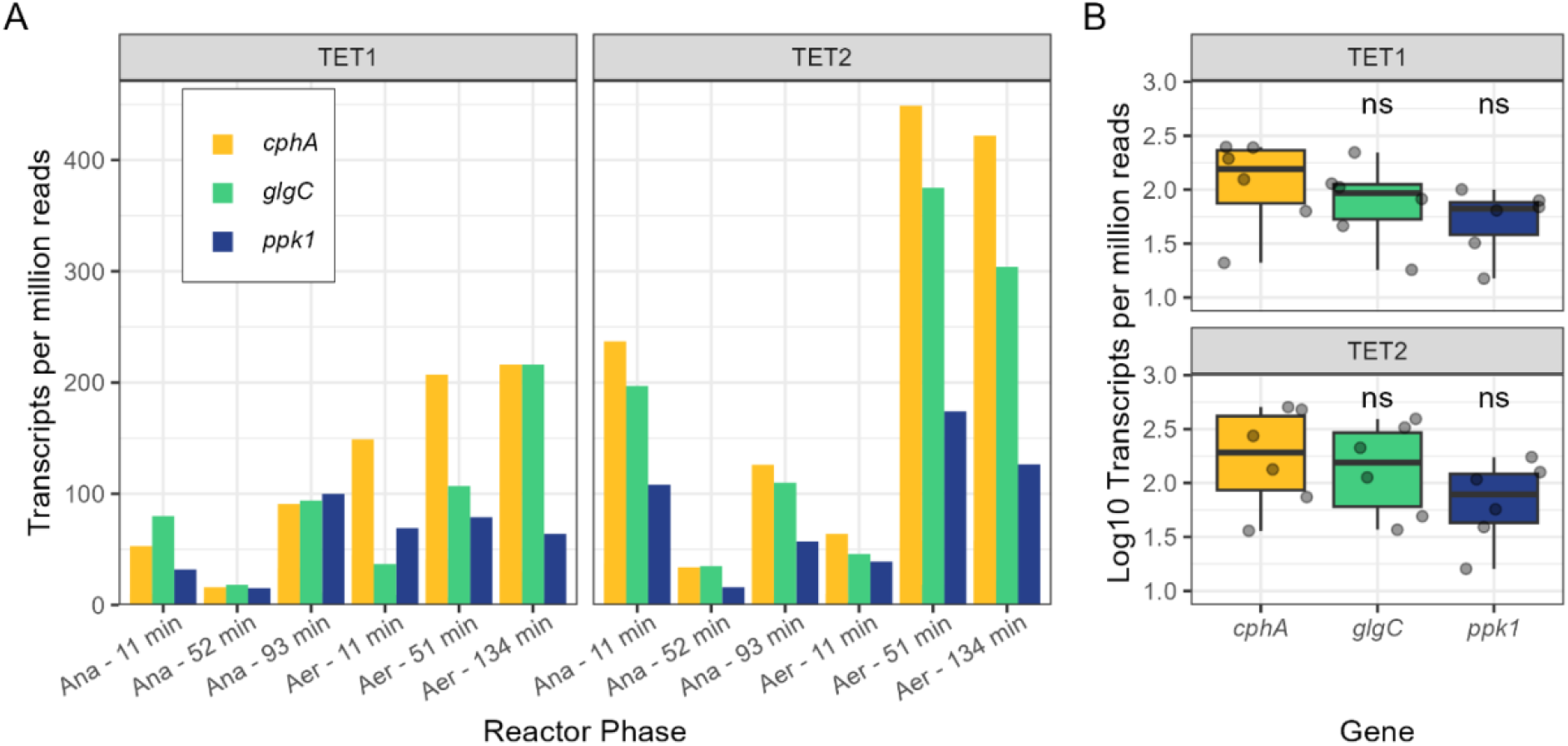
Gene expression profiles of *cphA* (expression levels of both copies summed), *glgC*, and *ppk* in *Tetrasphaera* MAGs plotted by the reactor phase (Ana=anaerobic, Aer=aerobic) (A) and summarized per MAG (B) of TET1 and TET2 from McDaniel et al., 2022. Significance levels in panel B are based on the Wilcox rank sum test, where *glgC*, and *ppk* are compared against *cphA* in each subpanel.

The expression of *cphA* in PAO *Ca*. Accumulibacter and *Tetrasphaera* is promising for future applications of combined P and N accumulation. In particular, the location of the *cphA* gene in *Tetrasphaera* near other key carbon cycling genes, such *glgA* and *glgC* for glycogen synthesis, increases the likelihood that cyanophycin is an actively used biopolymer for these bacteria. We also observed gene expression of *cphB* in the *Tetrasphaera* MAGs (Figure S6), further increasing the likelihood that cyanophycin is actively produced and used by these PAO. Further analyses of cyanophycin pathway activity in response to operational variables and measurements of cyanophycin in activated sludge biomass would improve our certainty of the possibilities for simultaneous N and P bioconcentration in PAO.

## 4. Conclusion and Outlook

In this study, we examined the prevalence of cyanophycin synthesis genes in wastewater bioprocess microbiomes. We observed an unexpectedly high prevalence of the *cphA* gene across a wide phylogenetic spectrum of common bacterial taxa in wastewater bioprocesses. The potential of cyanophycin accumulation in tandem with other nutrient cycling processes is promising given the seeming ubiquity of cyanophycin synthetase in common PAO *Ca*. Accumulibacter, *Tetrasphaera*, and *Dechloromonas* and nitrifiers *Nitrosomonas* and *Nitrosospira*. We also used metaetranscriptomic profiling to determine whether *cphA* genes were expressed by PAO under typical operating conditions, and we found expression levels similar to other important P and carbon cycling genes. Overall, the presence of cyanophycin synthetase in nutrient cycling taxa suggests that cyanophycin cycling may already be occurring in existing biological nutrient removal processes.

Further research will expand on the findings of this work. First, the feasibility of integrating cyanophycin accumulation into existing nutrient removal processes will largely depend on the ability to modulate cyanophycin production in concert with other desired functions, particularly P accumulation. Although we found that PAO harbor and express *cphA*, it is not clear whether cyanophycin accumulation occurs simultaneously with P accumulation. Second, our fundamental understanding of cyanophycin accumulation by wastewater bioprocess taxa will improve with further examination of the *cphA* gene cluster as a possible mobile genetic element. While mobile genetic elements are intensely studied as a means of transferring antibiotic resistance genes in wastewater-associated microbiomes, their role in transferring nutrient cycling genes is less clear. Overall, our findings provide evidence that cyanophycin accumulation is a widespread metabolic pathway in nutrient removal bioprocesses and opens possibilities for accelerating nutrient recovery from wastewater through N bioconcentration.

