## Supplemental information for "Meta-omic profiling reveals ubiquity of genes encoding for the high value nitrogen rich biopolymer cyanophycin in activated sludge microbiomes"

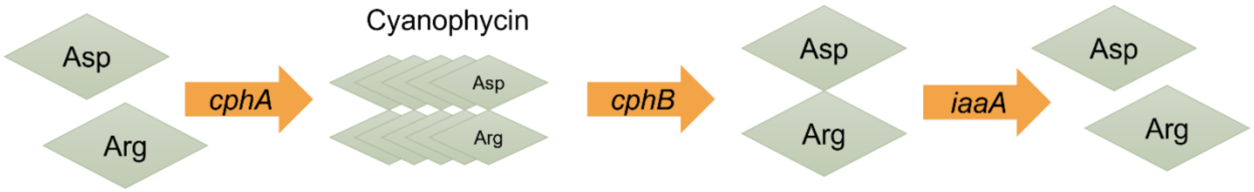


**Figure 1. Cyanophycin metabolic pathway.**

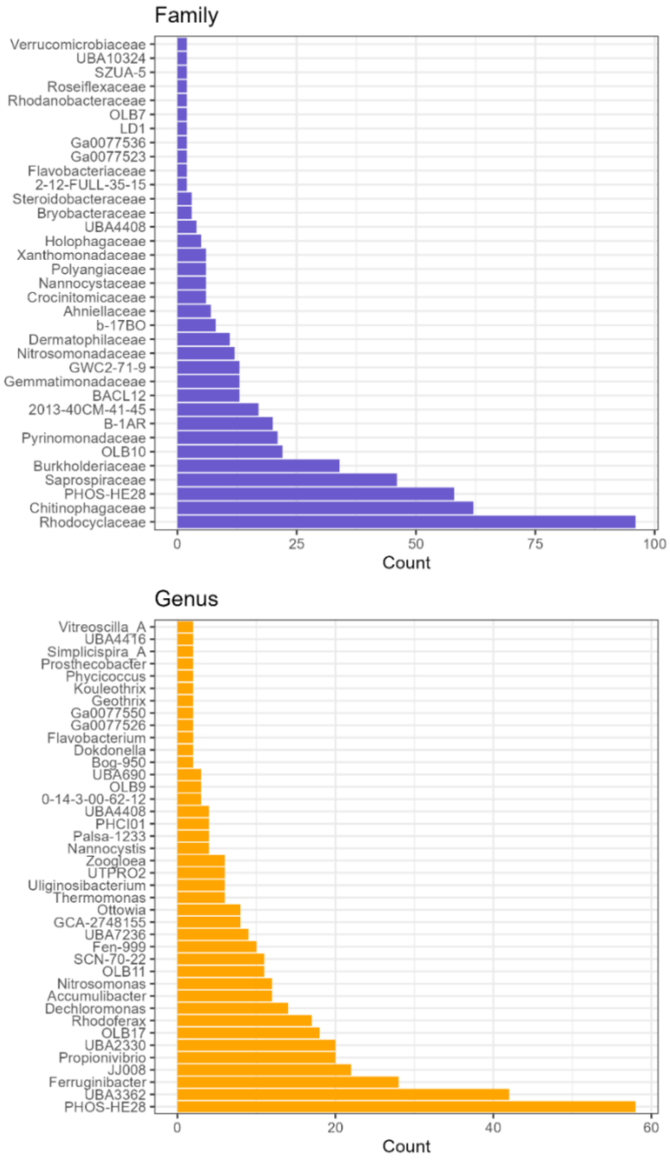


**Figure 2. Family (top) and genus (bottom) classifications of MAGs from the Singleton et al., 2021 dataset that contained a *cphA* gene. Data shown in this figure are for family or genus classifications of two or more MAGs. Unknown taxonomic classifications are not shown.**

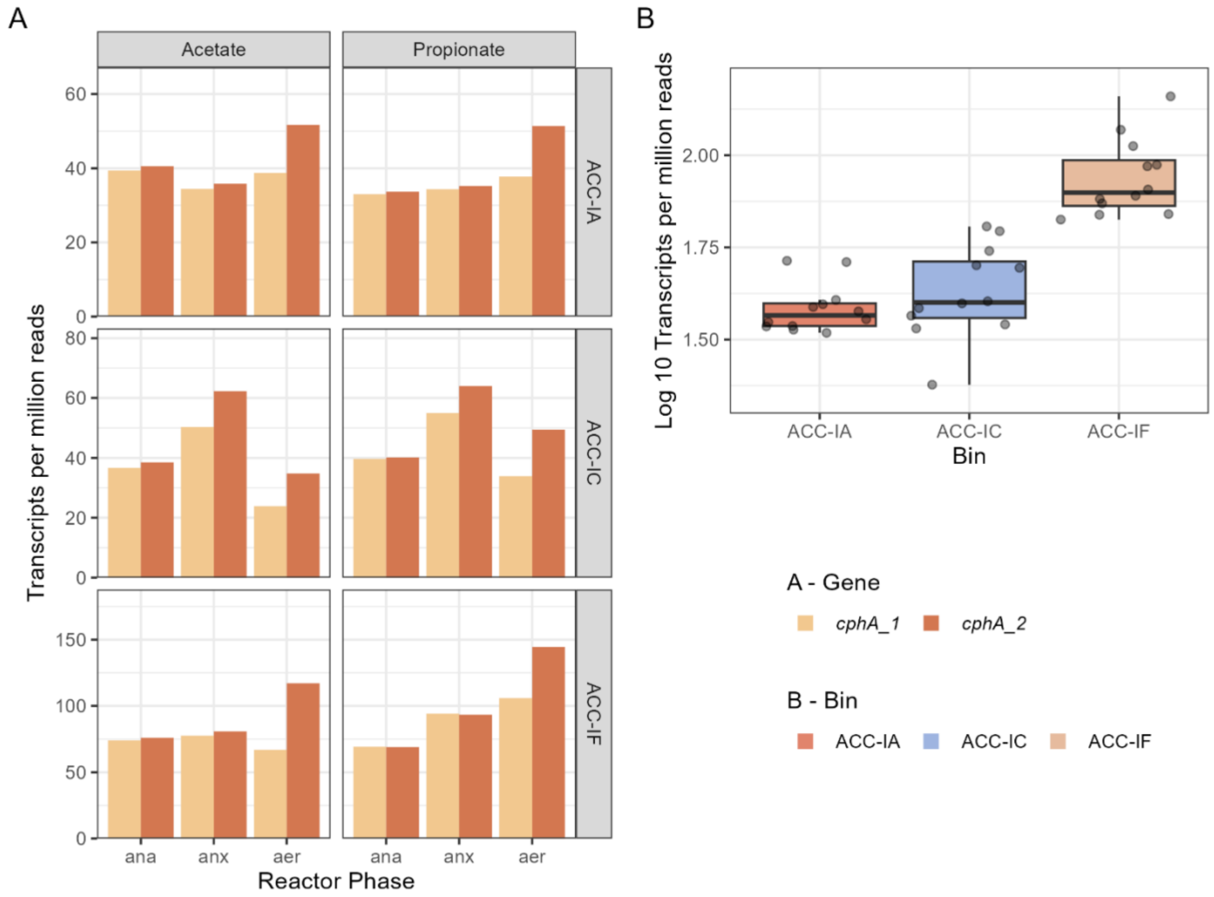


**Figure 3. Gene expression profiles of *cphA* in *Ca.* Accumulibacter MAGs from Gao et al., 2019 and Wang et al., 2021 by *cphA* gene copy, reactor phase and carbon source (A) and both gene copies summarized by bin (B).**

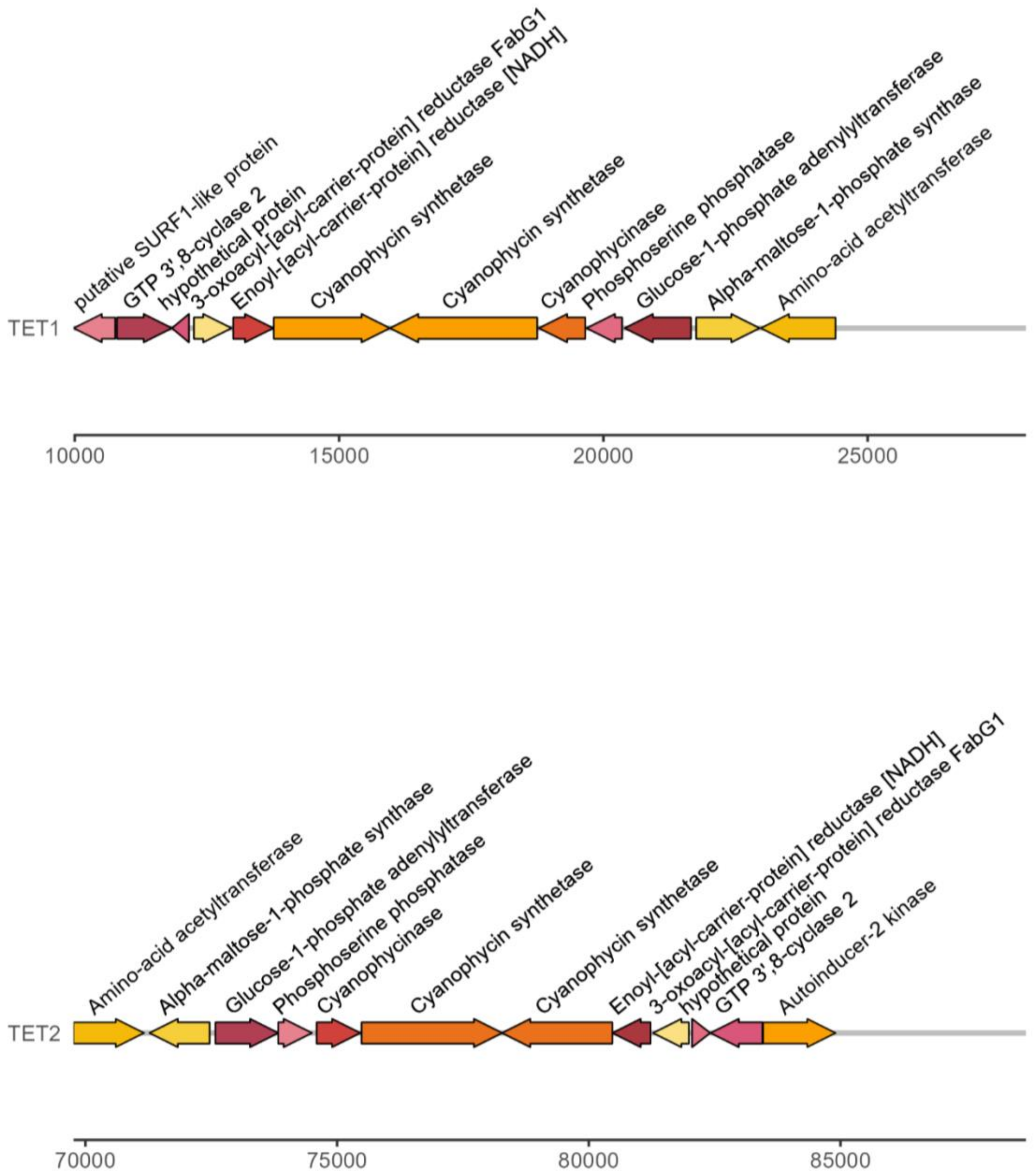


**Figure 4. Gene map of TET1 and TET2 from McDaniel et al., 2022 around the *cphA* gene**

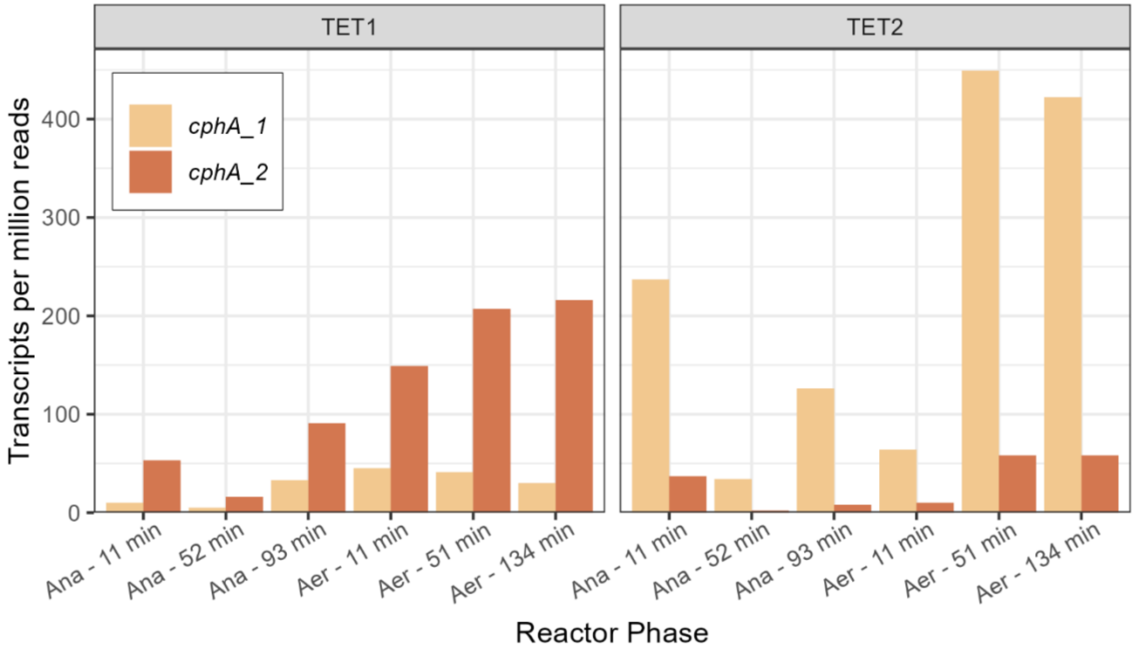


**Figure 5. Gene expression profiles of *cphA* in *Tetrasphaera* MAGs from McDaniel et al. 2022 by bin and *cphA* gene copy. Ana = anaerobic and Aer = aerobic.**

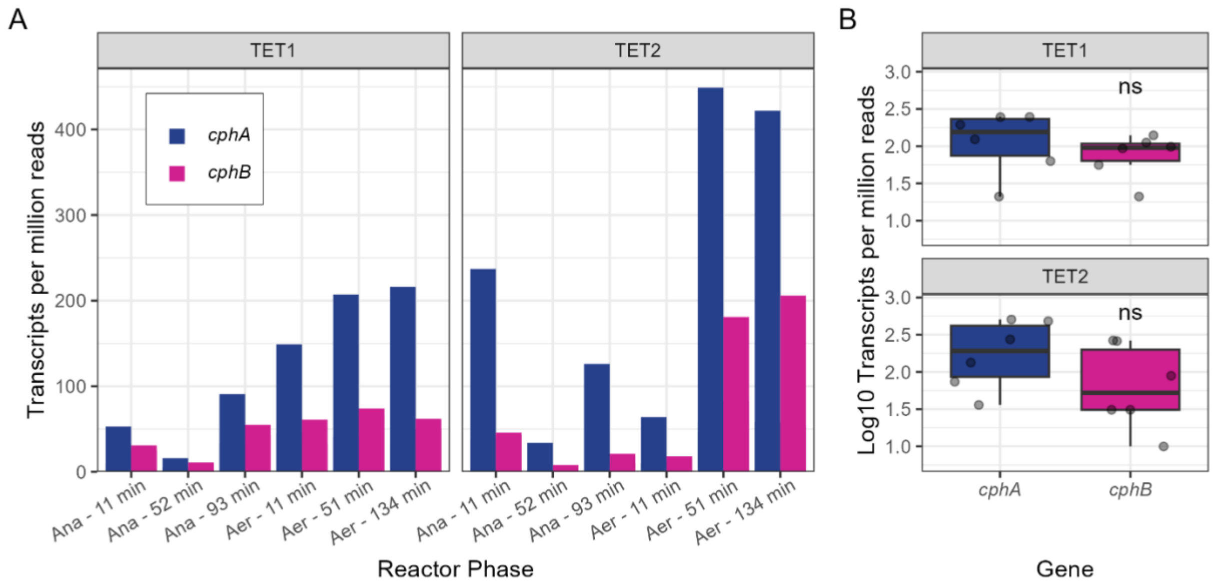


**Figure 6. Gene expression profiles of *cphA* and *cphB* in *Tetrasphaera* MAGs from McDaniel et al. 2022. Ana = anaerobic and Aer = aerobic.**
